# Cryo-EM structure of the endothelin-1-ET_B_-G_i_ complex

**DOI:** 10.1101/2023.01.12.523720

**Authors:** Fumiya K. Sano, Hiroaki Akasaka, Wataru Shihoya, Osamu Nureki

**Author notes:** Correspondence to (W.S.); (O.N.).

## Abstract

The endothelin ET_B_ receptor is a promiscuous G-protein coupled receptor, activated by vasoactive peptide endothelins. ET_B_ signaling induces reactive astrocytes in the brain and vasorelaxation in vascular smooth muscle, and thus ET_B_ agonists are expected to be utilized for neuroprotection and improved anti-tumor drug delivery. Here, we report a cryo-electron microscopy structure of the endothelin-1-ET_B_-G_i_ complex at 2.8-Å resolution, with complex assembly stabilized by a newly established method. Comparisons with the inactive ET_B_ receptor structures revealed how endothelin-1 activates the ET_B_ receptor. The NPxxY motif, which is essential for G-protein activation, is not conserved in ET_B_, resulting in a unique structural change upon G-protein activation. As Compared with other GPCR-G-protein complexes, ET_B_ binds G_i_ at the shallowest position, thus expanding the diversity of G-protein binding. This structural information will facilitate the elucidation of G-protein activation and the rational design of ET_B_-agonists.

## Introduction

Endothelins are 21-amino-acid vasoconstricting peptides produced primarily in the endothelium^1^, and play a key role in vascular homeostasis. Among the three endothelin isopeptides (ET-1 to 3), ET-1 was first discovered as a potent vasoconstrictor^2,3^. ET-1 transmits signals through two receptor subtypes, the ET_A_ and ET_B_ receptors, which belong to the class A G-protein-coupled receptors (GPCRs)^4,5^. ET-1 has a strong vasoconstrictor effect through the activation of ET_A_, while ET_B_ activation has different functions^3,6,7^. The ET_B_ receptor couples to both of G_i_ and G_q_^8^.and is reported to be a promiscuous GPCR, which can activate all G-protein subtypes^9^. In endothelium, the ET_B_-receptor-mediated G_q_ signaling induces nitric oxide-mediated vasorelaxation. In addition, the astrocytic ET_B_-receptor-mediated G_i_ signal is reportedly involved in an ET-induced reduction in intercellular communication through gap junctions^10^. Moreover, the activation of the ET_B_-receptor-mediated Rho signal in astrocytes involves cytoskeletal reorganization and cell-adhesion-dependent proliferation^11^. These signal mechanisms triggered by ET_B_ receptor are thought to underlie the induction of reactive astrocytes to promote neuroprotection^12^. Thus, ET_B_ selective agonists have been studied as vasodilator drugs for the improvement of tumor drug delivery, as well as for the treatment of brain damage^3^. IRL1620, a truncated peptide analog of ET-1, is the smallest agonist that can selectively stimulate the ET_B_ receptor, which is used in clinical studies by injection^13^. Currently, small-molecule ET_B_-selective agonists have not been developed, limiting the drug development targeting the ET_B_ receptor.

To date, eight crystal structures of the human ET_B_ receptor have been reported, elucidating the structure-activity relationships of the peptide agonists and small-molecule clinical antagonists^14–18^. Nevertheless, the detailed activation mechanism has remained elusive because the crystallization constructs contain T4 lysozyme in the intracellular loop (ICL) 3 and thermostabilizing mutations^19^, which fix the inactive state on the intracellular side of the receptor. Moreover, the conserved N^7.49^P^7.50^xxY^7.53^ motif (superscripts indicate Ballesteros–Weinstein numbers^20^) essential for G-protein activation^21^ is altered to N^7.49^P^7.50^xxL^7.53^Y^7.54^ in the wild-type ET_B_ receptor. Thus, little is known about ET_B_-mediated G-protein activation. Here, we report the 2.8-Å resolution cryo-electron microscopy (cryo-EM) structure of the human ET_B_-G_i_ signaling complex bound to ET-1. Close examination of the ET_B_ structure has revealed the unique mechanisms of receptor activation and G-protein coupling.

## Results

### Development of Fusion-G system for structural determination

For the cryo-EM analysis, we initially used the thermostabilized receptor ET_B_-Y5, which contains five thermostabilizing mutations^19^. However, the purified ET_B_-Y5 could not form a stable complex with the G_i_ trimer, because the mutations reportedly stabilize the inactive conformation. Thus, we adopted the wild-type ET_B_ for the structural study. To purify the stable GPCR-G protein complex, we developed a “Fusion-G system” (Figure 1A) that combines two techniques for complex stabilization. One is a NanoBiT tethering strategy^22,23^, in which the large part of NanoBiT (LgBiT) was fused to the C-terminus of the receptor, and a renovated 13-amino acid peptide of NanoBiT (HiBiT) was fused to the C-terminus of Gβ via the GS linker. Hibit has potent affinity for LgBiT (*Kd*=700 pM), and thus provides an additional linkage to stabilize the interface between H8 of the receptor and the Gβ subunit of the G protein. This strategy has been successfully used in solving several GPCR/G-protein complex structures^22,24^. The other is a 3-in-1 vector for G-protein expression, in which the Gα subunit is fused to the C-terminus of the Gγ subunit^25,26^ (Figure 1A). The resulting pFastBac-Dual vector can produce a virus that expresses the G-protein trimer. Moreover, the protease-cleavable green fluorescent protein (EGFP) is connected to the C-terminus of the receptor-LgBiT fusion, allowing analysis of complex formation by fluorescence-detection size-exclusion chromatography (FSEC)^27^. Using this system, we confirmed the complex formation of LPA_1_ and S1P_5_ (Figure 1–figure supplement 1), whose structures in complex with the G_i_ trimer were previously reported^28–31^.

**Figure 1.**
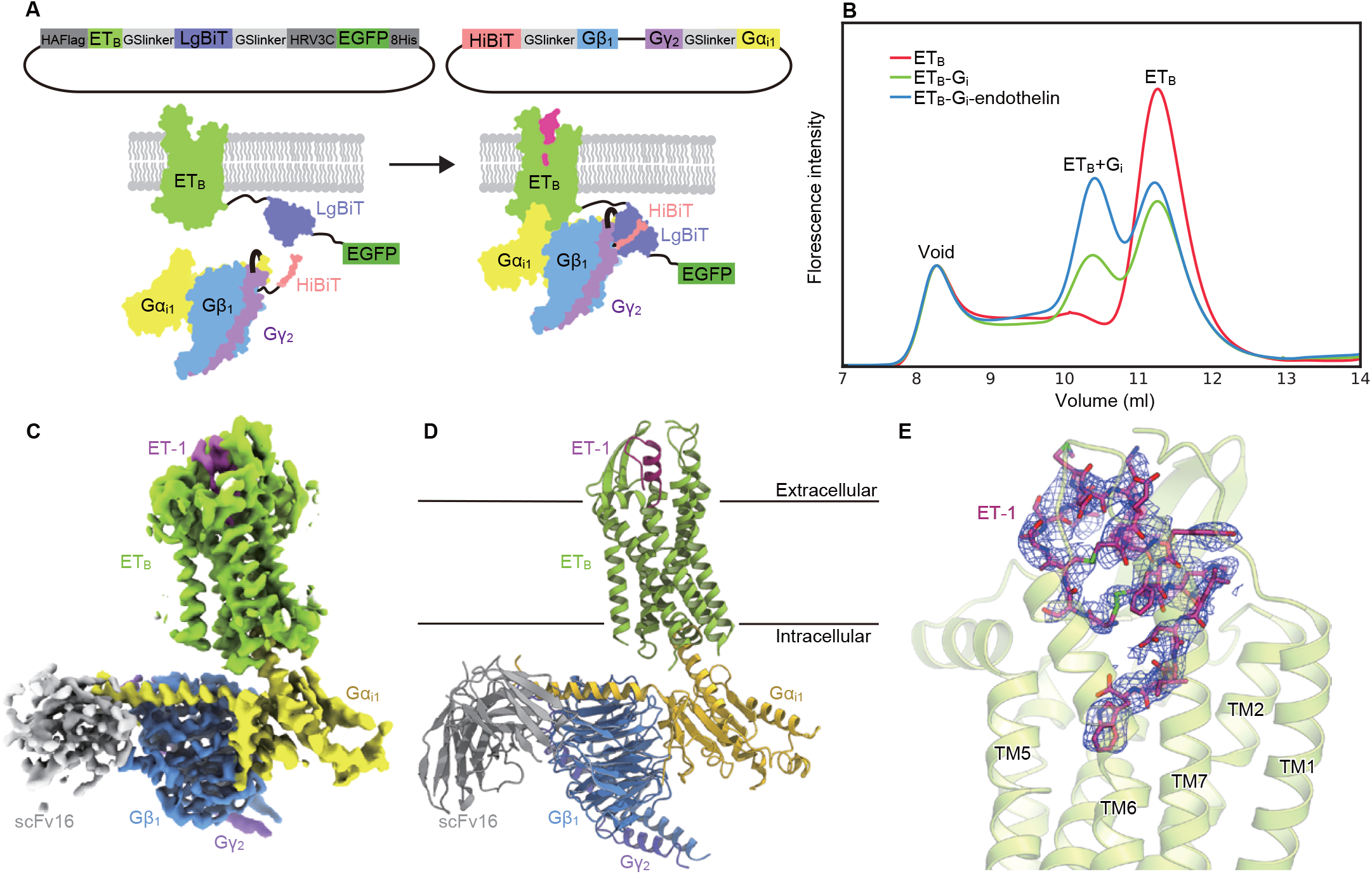
Overall structure of the ET-1-ET_B_-G_i_ signaling complex. (A) Schematic representations of the fusion-G system. (B) FSEC analysis of complex formation by the ET_B_ receptor. (C) Sharpened cryo-EM map with variously colored densities. Overall refined map and focused refined map were combined by phenix.combine_focused_maps. (D) Structure of the complex determined after refinement in the cryo-EM map, shown as a ribbon representation. (E) Density focused on ET-1.

We cloned the full-length human ET_B_ receptor into the LgBiT vector. Using the fusion-G-system, we confirmed the formation of the ET_B_-G_i_ complex (Figure 1B). The co-expressed cells from a 300 ml culture were solubilized and purified by Flag affinity chromatography. After an incubation with scFv16, the complex was purified by size exclusion chromatography (Figure 1–figure supplement 1). The structure of the purified complex was determined by single-particle cryo-EM analysis with an overall resolution of 2.8 Å (Figure 1C, D, Figure 1–figure supplement 2, Table 1). No density corresponding to NanoBiT was observed in the 2D class averages and reconstructed 3D density map, suggesting that the fusion G system minimally affects the complex structure. We performed a refinement with a mask on the receptor and obtained the receptor structure with a nominal resolution of 3.1 Å (Figure 1C, D, Figure 1–figure supplement 2, Table 1). The agonist ET-1 is well resolved (Figure 1E).

**Table 1.** Cryo-EM data collection, refinement and validation statistics.

| Data collection | ET<sub>B</sub>-G<sub>i</sub> (overall) | ET<sub>B</sub>-G<sub>i</sub> (receptor focused) |
| --- | --- | --- |
| Microscope | Titan Krios (Thermo Fisher Scientific) |  |
| Voltage (keV) | 300 |  |
| Electron exposure (e<sup>-</sup>/Å<sup>2</sup>) | 49.965 |  |
| Detector | Gatan K3 summit camera (Gatan) |  |
| Magnification | ×105,000 |  |
| Defocus range (μm) | -0.8–1.6 |  |
| Pixel size (Å/pix) | 0.83 |  |
| Number of movies | 10,408 |  |
| Symmetry | C1 |  |
| Picked particles | 3,863,134 |  |
| Final particles | 260,085 |  |
| Map resolution (Å) | 2.8 | 3.13 |
| FSC threshold | 0.143 |  |
| Model refinement |  |  |
| Atoms | 9,082 | 2,533 |
| R.m.s. deviations for ideal |  |  |
| Bond lengths (Å) | 0.0030 | 0.0027 |
| Bond angles (°) | 0.71 | 0.63 |
| Validation |  |  |
| Clashscore | 14.99 | 9.14 |
| Rotamers (%) | 0.2 | 0.35 |
| Ramachandran plot |  |  |
| Favored (%) | 95.95 | 98.71 |
| Allowed (%) | 3.96 | 1.29 |
| Outlier (%) | 0.09 | 0 |

### Receptor conformational changes upon G_i_ activation

The extracellular half of the receptor superimposes well on the ET-1-bound crystal structure, and ET-1 forms similar tight interactions with the receptor (Figure 2A). Previous crystallographic analyses have suggested that ET-1 binding induces the downward movements of N378^7.45^ and W336^6.48^ in the C^6.47^W^6.48^xP^6.50^ motif at the bottom of the ligand binding pocket, resulting in the downward rotation of F332^6.44^ in the P^5.50^I^3.40^F^6.44^ motif, and leading to the intracellular opening^14–18^. In the ET_B_-G_i_ complex, the downward movements of the residues are larger than those in the ET-1-bound crystal structure (Figure 2B), resulting in the outward displacement of the intracellular portion of transmembrane helix (TM) 6 by 6.8 Å as compared to the apo state (Figure 2C), and by 5.1 Å as compared to the ET-1-bound crystal structure (Figure 2D). The observed TM6 opening is smaller than those in other G_i_-coupled receptors (e.g., μOR: 10 Å, CB_1_: 11.6 Å, and S1P_1_: 9 Å)^30,32,33^.

**Figure 2.**
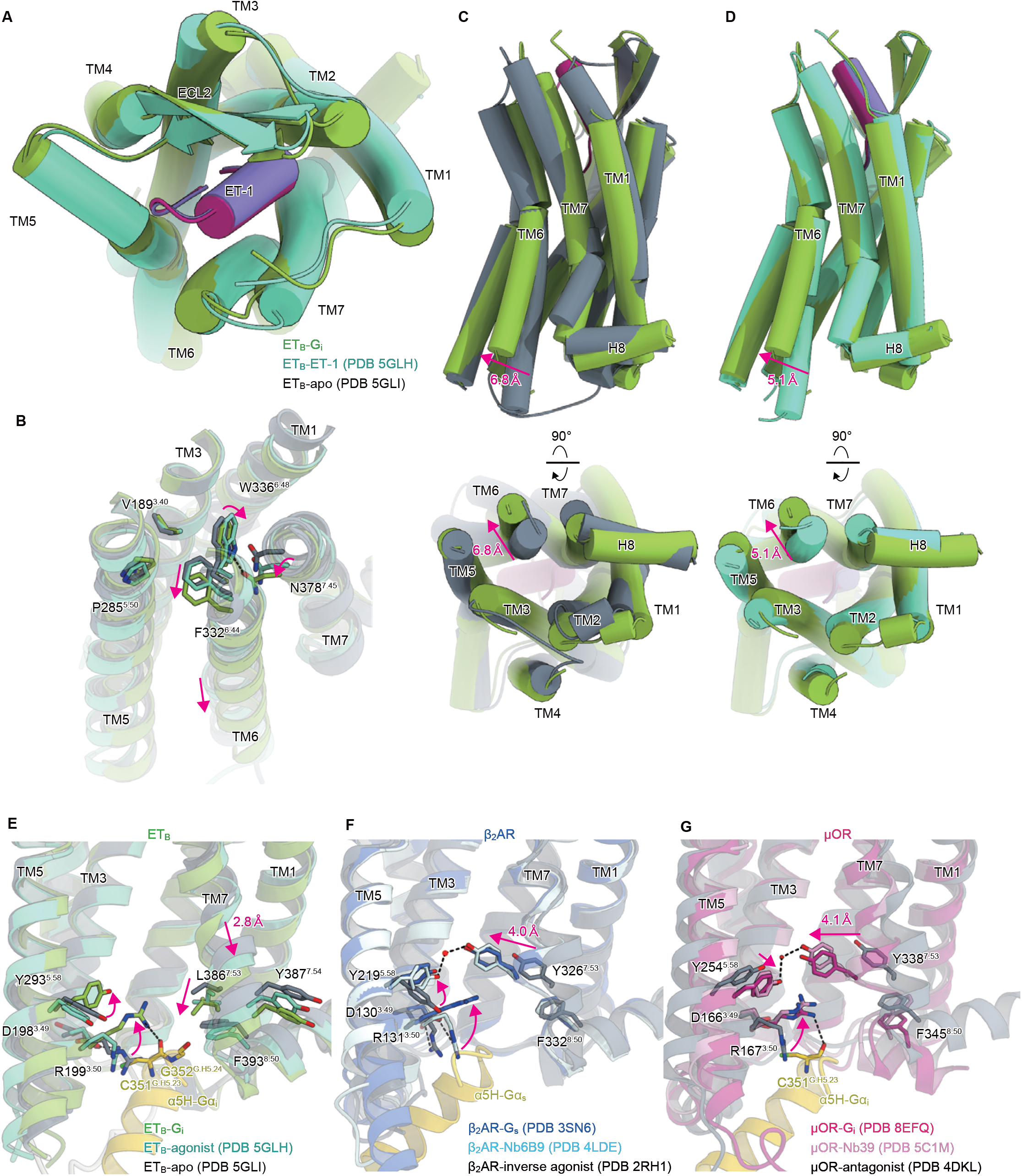
Structural changes upon G-protein activation. (A) Superimposition of the ET-1-bound receptor in the crystal and cryo-EM structures. (B) Superimposition of the ET_B_ structures, focused on the receptor core. (C, D) Superimpositions of the G_i_-complexed ET_B_ structure with the ET-1-bound crystal structure (C) and apo structure (D). (E-G) D^3.49^R^3.50^Y^3.51^ and N^7.49^P^7.50^xxY^7.53^ motifs in ET_B_ (E), β_2_AR (F), and μOR (G). Black dashed lines indicate hydrogen bonds.

In most class A GPCRs, the highly conserved D^3.49^R^3.50^Y^3.51^ and N^7.49^P^7.50^xxY^7.53^ motifs exist on the intracellular side, where they play an essential role in G-protein coupling^21^. The D^3.49^R^3.50^Y^3.51^ motif is conserved in the ET_B_ receptor. Upon receptor activation, the ionic lock between D198^3.49^ and R199^3.50^ is broken, and R199^3.50^ becomes oriented toward the intracellular cavity (Figure 2E), as in other GPCRs^32,34–38^ (Figure 2F, G). In contrast, the N^7.49^P^7.50^xxY^7.53^ motif is altered to N^7.49^P^7.50^xxL^7.53^Y^7.54^, in which Y^7.53^ is replaced by L386^7.53^. In most class A GPCRs, receptor activation disrupts the stacking interaction between Y^7.53^ and F^8.50 39^. Y^7.53^ moves inwardly and forms a water-mediated hydrogen bond with the highly conserved Y^5.58^ (Figure 2F, G)^35,38,40^. The mutations of the tyrosines greatly reduce activation of G-protein^41,42^, indicating that Y^5.58^-Y^7.53^ interaction stabilizes the active conformation of the receptor. Along with the motion, the intracellular portion of TM7 is displaced by about 4 Å. In the ET_B_-G_i_ complex, as the hydrophobic residue L386^7.53^ cannot form such a polar interaction, TM7 is not displaced inwardly (Figure 2C). Nevertheless, the stacking interaction with F393^8.50^ is similarly disrupted. Moreover, the intracellular portion of TM7 is displaced downward by 2.8 Å. As expected, Y^7.54^ is directed towards the membrane plane and its rotamer does not change upon receptor activation (Figure 2E). The replacement of Y^7.53^ with L386^7.53^ affects the movement of TM7 upon ET_B_ receptor activation, thus distinguishing it from other GPCRs.

### Shallow G_i_ coupling

These conformational changes create an intracellular cavity for G-protein recognition (Figure 3A, Figure 3–figure supplement 1). The cavity closely contacts the C-terminal α5-helix of Gα_i_, which is the primary determinant for G-protein coupling. Specifically, R199^3.50^ forms a hydrogen bond with the backbone carbonyl of C351^G.H5.23^ (superscript indicates the common Gα numbering [CGN] system^42^), as commonly observed in other GPCR-G_i_ complexes^30,32,33,43^. Moreover, the C-terminal carboxylate of α5-helix forms electrostatic interactions with the backbone nitrogen atom of R391^8.48^. There are several other hydrogen-bonding interactions between the α5-helix and the receptor (Figure 3A). Although most of ICL2 is disordered, W206^ICL2^ fits into hydrophobic pocket formed by L194^G.S3.01^, F336^G.H5.08^, T340^G.H5.12^, I343^G.H5.15^, and I344^G.H5.16^ in the Gα_i_ subunit (Figure 3–figure supplement 1), as in the other GPCR-G_i_ complexes^30,33^.

**Figure 3.**
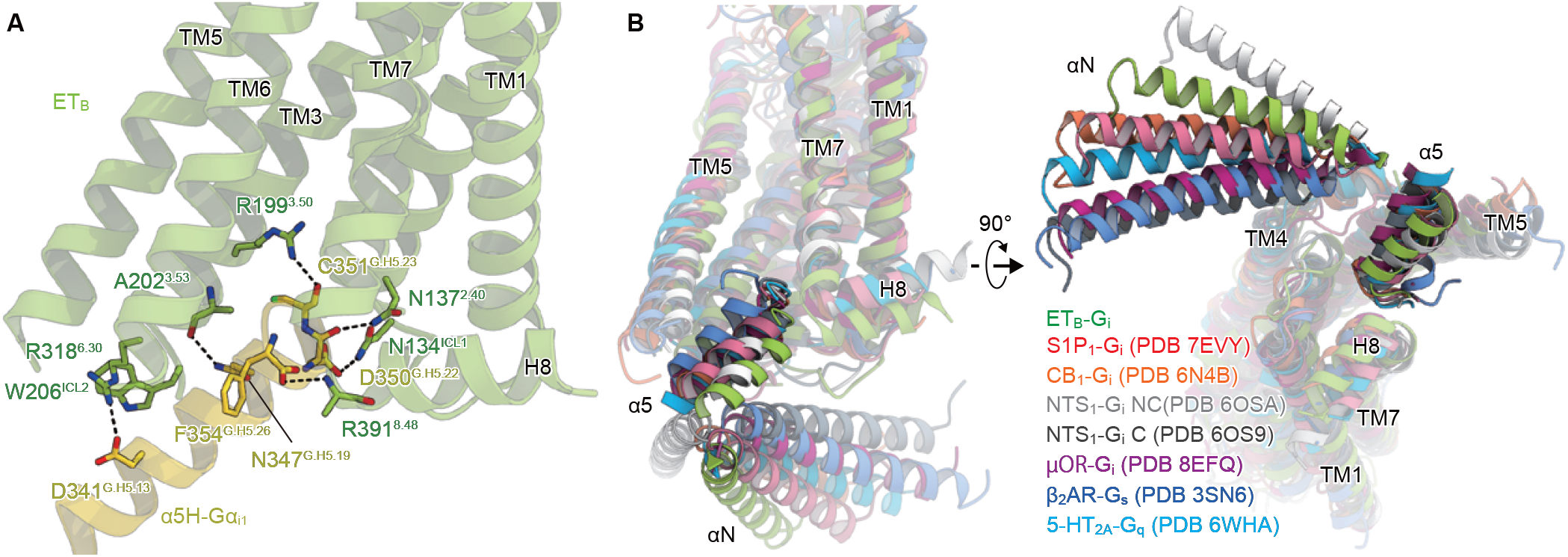
Comparison of the G_i_ binding modes. (A) Hydrogen-bonding interactions between ET_B_ and the α5-helix, indicated by black dashed lines. (B) Comparison of the Gα positions in the GPCR-G-protein complexes. The structures are superimposed on the receptor structure of the NTSR1-C state.

Since the intracellular position of TM7 is displaced downward, L386^7.53^ directly forms a hydrophobic contact with G352^G.H5.25^. Moreover, TM7 and H8 extensively interact with the α5-helix (Figure 3–figure supplement 2). These interactions are not observed in other GPCR-G_i_ complexes (Figure 3–figure supplement 2) ^30,32,33,43^, and are unique to the ET_B_-G_i_ interface. This feature in TM7 allows the α5-helix of ET_B_-G_i_ to become located in the shallowest position relative to the receptor, among the Gs, G_i_, and G_q_-coupled GPCR structures^25,30,32–34,43^ (Figure 3B). However, the Gα_i_ structure adopts a nucleic acid-free state, as in the μOR-G_i_ complex (Figure 3–figure supplement 1)^32,44^. This comparison indicates that the G-protein adopts active states, although the α5-helix is located in the shallowest position.

## Discussion

In this study, we established the Fusion-G system, which enables the efficient expression and purification of the GPCR-G-protein complex. In addition to this system, the ability to monitor the fluorescence of GFP fused in the C-terminus of the receptor makes it easier to optimize the expression and purification conditions. The ET_B_-G_i_ structure determined by this strategy filled in the last piece of the puzzle, and our understanding of the mechanism of the receptor activation has been significantly deepened. The full-active structure of the bottom of the ligand binding pocket will pave the way for the development of small molecule ET_B_ agonists (Figure 2B).

Previous structural studies have shown that the docking of the α5-helix to the receptor cavity destabilizes the nucleic acid binding site at the root of the α5-helix (Figure 3–figure supplement 1)^32,34,45^, thus promoting the GDP/GTP exchange reaction. Whereas the Gα_s_ binding is essentially similar in all of the complexes, the Gα_i_ binding is variable, with different Gα rotations relative to the receptor^46^. Nevertheless, the depth of the docking toward the receptor was constant (Figure 3B). Notably, a previous time-resolved mass spectroscopic analysis of β_2_AR suggested that the initial engagement of the α5-helix and ICL2 is sufficient for G-protein activation, and the deep docking of the α5-helix seen in previous structures is not necessarily involved in G-protein activation^47^. Supporting this notion, the α5-helix of ET_B_-G_i_ is located in the shallowest position (Figure 3B). Despite the absence of the critical Y^7.53^ in the conserved N^7.49^P^7.50^xxY^7.53^ motif, ET_B_ distinctly binds and activates G_i_. Such information is essential for understanding the mechanism of G_i_ activation by GPCRs.

## Materials and Methods

### Constructs

The full-length human ET_B_ gene was subcloned into the pFastBac vector with a HA-signal peptide sequence on its N-terminus and the LgBiT fused to its C-terminus followed by a 3C protease site and EGFP-His8 tag. A 15 amino sequence of GGSGGGGSGGSSSGG was inserted into both N-terminal and C-terminal sides of LgBiT. The Flag epitope tag (DYKDDDDK) was introduced between G57 and L66. The native signal peptide was replaced with the haemagglutinin signal peptide. Rat Gβ_1_ and bovine Gγ_2_ were subcloned into the pFastBac Dual vector, as described previously^48^. In detail, rat Gβ_1_ was cloned with a C-terminal HiBiT connected with a 15 amino sequence of GGSGGGGSGGSSSGG. Moreover, human Gα_i1_ was subcloned into the C-terminus of the bovine Gγ_2_ with a 9 amino sequence GSAGSAGSA linker. The resulting pFastBac dual vector can express the G_i_ trimer.

### Complex formation and FSEC analysis

Bacmid preparation and virus production were performed according to the Bac-to-Bac baculovirus system manual (G_i_bco, Invitrogen). *Spodoptera frugiperda* Sf9 cells at a density of 3 × 10^6^ cells/ml were co-infected with baculoviruses encoding receptor and G_i_ trimer at the ratio of 1:1. For the expression of the receptor alone, the baculovirus encoding receptor was only used. Cells were harvested 48 h after infection. 1 ml cell pellets were solubilized in 200 μl buffer, containing 20 mM Tris-HCl, pH 8.0, 150 mM NaCl, 1% n-dodecyl-β-D-maltoside (DDM) (Calbiochem), 0.2 % cholesteryl hemisuccinate (CHS) (Merck) and rotated for 1 h at 4 °C.

For the complex formation with the agonist, cell pellets were resuspended in 20 mM Tris-HCl, pH 8.0, 100 mM NaCl, and 10% Glycerol, and homogenized by douncing ~20 times. Apyrase was added to the lysis at a final concentration of 25 mU/ml. Each agonist was added at a final concentration of 10 μM. The homogenate was incubated at room temperature for 1 h with flipping. Then, DDM and CHS were added to a final concentration of 1% and 0.2 %, respectively for 1 h at 4 °C.

The supernatants were separated from the insoluble material by ultracentrifugation at 100,000*g* for 20 min. A fraction of the resulting supernatant (10 μl) was loaded onto a Superdex 200 increase 10/300 column in 20 mM Tris-HCl, pH 8.0, 150 mM NaCl, and 0.03% DDM, and runed at the flow rate of 0.5 ml/min. The eluent was detected by a fluorometer with the excitation wavelength (480 nm) and emission wavelength settings (512 nm).

### ET-1–ET_B_–G_i_ complex formation and purification

For expression, 300 ml of the Sf9 cells at a density of 3 × 10^6^ cells/ml were co-infected with baculovirus encoding the ET_B_-LgBiT-EGFP and G_i_ trimer at the ratio of 1:1. Cells were harvested 48 h after infection. Cell pellets were resuspended in 20 mM Tris-HCl, pH 8.0, 100 mM NaCl, and 10% Glycerol, and homogenized by douncing ~30 times. Apyrase was added to the lysis at a final concentration of 25 mU/ml. ET-1 was added at a final concentration of 2 μM. The lysate was incubated at room temperature for 1 h with flipping. Then, the membrane fraction was collected by ultracentrifugation at 180,000*g* for 1h. The cell membrane was solubilized in buffer, containing 20 mM Tris-HCl, pH 8.0, 150 mM NaCl, 1% DDM, 0.2 % CHS, 10% glycerol, and 2 μM ET-1 for 1 h at 4 °C. The supernatant was separated from the insoluble material by ultracentrifugation at 180,000*g* for 30 min and then incubated with the Anti-DYKDDDDK G1 resin (Genscript) for 1 h. The resin was washed 20 column volumes of wash buffer containing 20 mM Tris-HCl, pH 8.0, 500 mM NaCl, 10% Glycerol, 0.1% Lauryl Maltose Neopentyl Glycol (LMNG) (Anatrace) and 0.01 % CHS. The complex was eluted by the wash buffer containing 0.15 mg ml^-1^ Flag peptide. The eluate was treated with 0.5 mg of HRV-3C protease (home made) and dialysed against buffer (20 mM Tris-HCl, pH 8.0, and 300 mM NaCl). Then, cleavaged GFP-His8 and HRV-3C protease were removed with Ni^+^-NTA resin. The flow through was incubated with the scFv16, prepared as described previously^46^. The complex was concentrated and loaded onto a Superdex 200 increase 10/300 column in 20 mM Tris-HCl, pH 8.0, 150 mM NaCl, 0.01% LMNG, 0.001% CHS, and 1 μM agonist. Peak fractions were concentrated to 8 mg/ml.

### Cryo-EM grid preparation and data acquisition

The purified complex was applied onto a freshly glow-discharged Quantifoil holey carbon grid (R1.2/1.3, Au, 300 mesh), and plunge-frozen in liquid ethane by using a Vitrobot Mark IV (FEI). Cryo-EM data collection was performed on a 300 kV Titan Krios G3i microscope (Thermo Fisher Scientific) equipped with a BioQuantum K3 imaging filter (Gatan) and a K3 direct electron detector (Gatan). In total, 10,408 movies were acquired with a calibrated pixel size of 0.83 Å pix^-1^ and with a defocus range of –0.8 to –1.6 μm, using the EPU software (Thermo Fisher’s single-particle data collection software). Each movie was acquired for 2.3 s and split into 48 frames, resulting in an accumulated exposure of about 49.965 e^−^ Å^−2^.

All acquired movies in super-resolution mode were binned by 2× and were dose-fractionated and subjected to beam-induced motion correction implemented in RELION 3.1^49^. The contrast transfer function (CTF) parameters were estimated using patch CTF estimation in cryoSPARC v3.3^50^. Particles were initially picked from a small fraction with the Blob picker and subjected to several rounds of two-dimensional (2D) classification in cryoSPARC. Selected particles were used for training of topaz model^51^. For the full dataset, 3,863,134 particles were picked and extracted with the pixel size of 3.32 Å, followed by 2D classification to remove ‘junk’ particles. A total of 1,442,243 particles were re-extracted with the pixel size of 1.16 Å and curated by three-dimensional (3D) classification without alignment in RELION. Finally, the 260,085 particles in the best class were reconstructed using non-uniform refinement, resulting in a 2.93 Å overall resolution reconstruction, with the gold standard Fourier Shell Correlation (FSC=0.143) criteria in cryoSPARC. Moreover, the 3D model was refined with a mask on the receptor. As a result, the local resolution of the receptor portion improved with a nominal resolution of 3.13 Å. The local resolution was estimated by cryoSPARC. The processing strategy is described in Figure 1–figure supplement 2.

### Model building and refinement

The quality of the density map was sufficient to build an atomic model. Previously reported high-resolution crystal structure of the ET-3 bound ET_B_ receptor (PDB 6IGK) and cryo-EM MT_1_-G_i_ structure (PDB 7DB6) were used as the initial models for the model building of receptor and G_i_ portions, respectively^17,46^. Initially, the models were fitted into the density map by jiggle fit using COOT^52^. Then, atomic models were readjusted into the density map using COOT and refined them using phenix.real_space_refine (v1.19) with the secondary structure restraints using phenix.secondary_structure_restraints^53,54^.

## Data availability

Cryo-EM density maps and atomic coordinates for the ET_B_-G_i_ complex have been deposited in the Electron Microscopy Data Bank and the PDB, under accession codes: XXXX.

## Author information

### Author details

Fumiya K. Sano

Department of Biological Sciences, Graduate School of Science, The University of Tokyo, 7-3-1 Hongo, Bunkyo-ku, Tokyo 113-0033, Japan.

Contribution: Conceptualization, Data curation, Formal analysis, Investigation, Methodology, Validation, Visualization, Writing – review and editing

Competing interests: No competing interests declared.

Hiroaki Akasaka

Department of Biological Sciences, Graduate School of Science, The University of Tokyo, 7-3-1 Hongo, Bunkyo-ku, Tokyo 113-0033, Japan.

Contribution Conceptualization, Data curation, Formal analysis, Investigation, Methodology, Validation, Visualization, Writing – review and editing

Competing interests: No competing interests declared.

Wataru Shihoya

Department of Biological Sciences, Graduate School of Science, The University of Tokyo, 7-3-1 Hongo, Bunkyo-ku, Tokyo 113-0033, Japan.

Contribution Conceptualization, Data curation, Formal analysis, Investigation, Methodology, Validation, Visualization, Project administration, Writing - original draft, Writing – review and editing

Competing interests No competing interests declared.

Osamu Nureki

Department of Biological Sciences, Graduate School of Science, The University of Tokyo, 7-3-1 Hongo, Bunkyo-ku, Tokyo 113-0033, Japan.

Contribution Conceptualization, Funding acquisition, Project administration, Supervision, Writing – review and editing

Competing interests: Co-founder and scientific advisor for Curreio.

## Acknowledgements

We thank K. Ogomori and C. Harada for technical assistance. This work was supported by grants from the Platform for Drug Discovery, Informatics and Structural Life Science by the Ministry of Education, Culture, Sports, Science and Technology (MEXT), and JSPS KAKENHI grants 21H05037 (O.N.), 22K19371 and 22H02751 (W.S.); ONO Medical Research Foundation (W.S.); The Kao Foundation for Arts and Sciences (W.S.); The Takeda Science Foundation (W.S.); The Uehara Memorial Foundation (W.S.); the Platform Project for Supporting Drug Discovery and Life Science Research (Basis for Supporting Innovative Drug Discovery and Life Science Research (BINDS)) from AMED, under grant numbers JP19am01011115 (support no. 1109, O.N.).

---

**Figure 1 – figure supplement 1.**
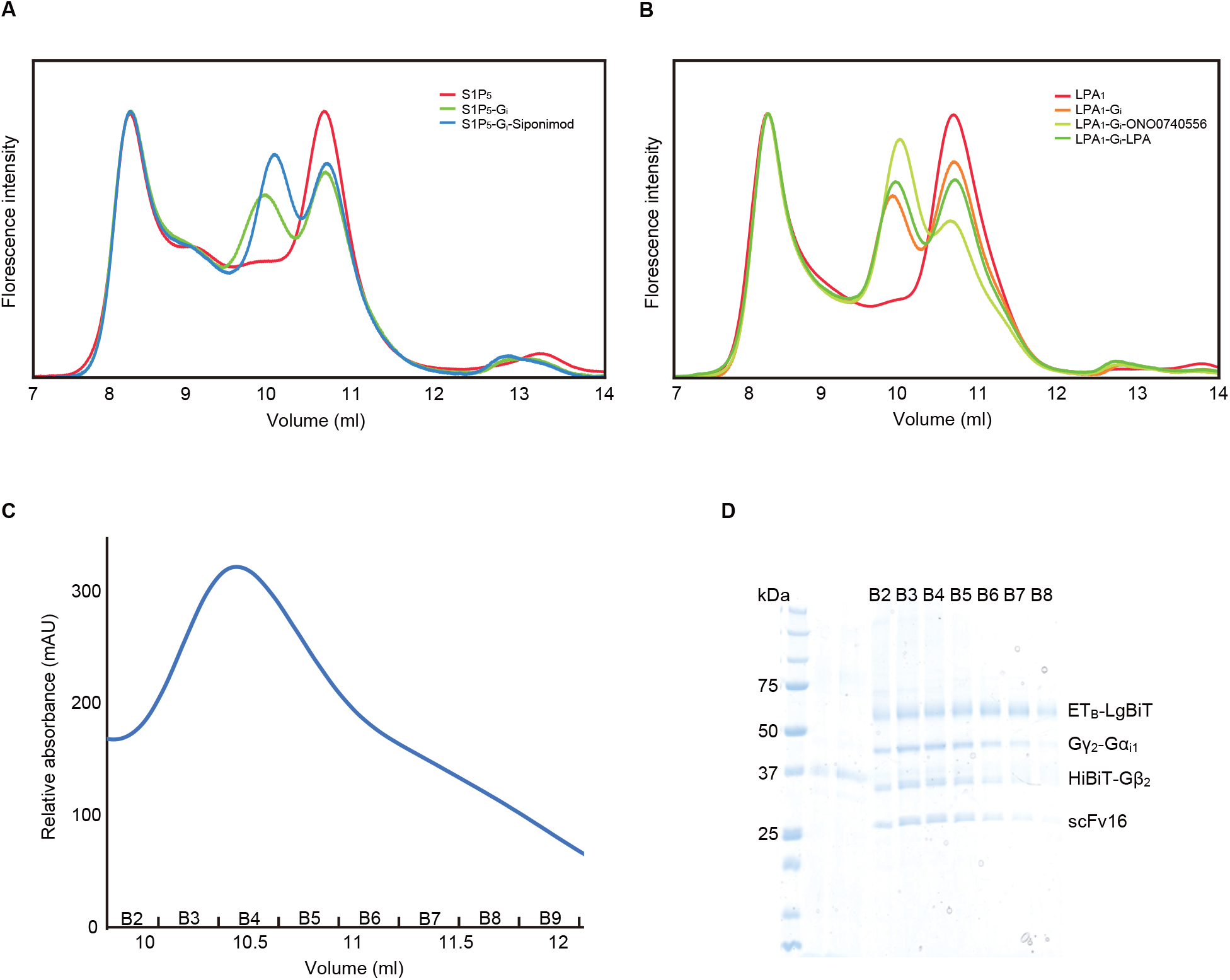
Fusion-G system. (A, B) FSEC analysis of the complex formation of LPA_1_ (A) and S1P_5_ (B). When co-expressed with the G_i_ trimer, the peak corresponding to the complex appears on the high molecular weight side of the receptor, reflecting the basal activity and the HiBiT-LgBiT binding, independent of the G-protein. When the co-expressed cells were treated with the agonists and apyrase, the complex peak of LPA_1_ and S1P_5_ became larger, corresponding to the GPCR-G protein complex. (C) Size-exclusion chromatography elution profiles of the ET_B_ -G_i_ complex. (D) SDS-PAGE analysis of the gel filtration fractions. Although the unbound receptor was present, the complex could be separated and purified effectively.

**Figure 1 – figure supplement 2.**
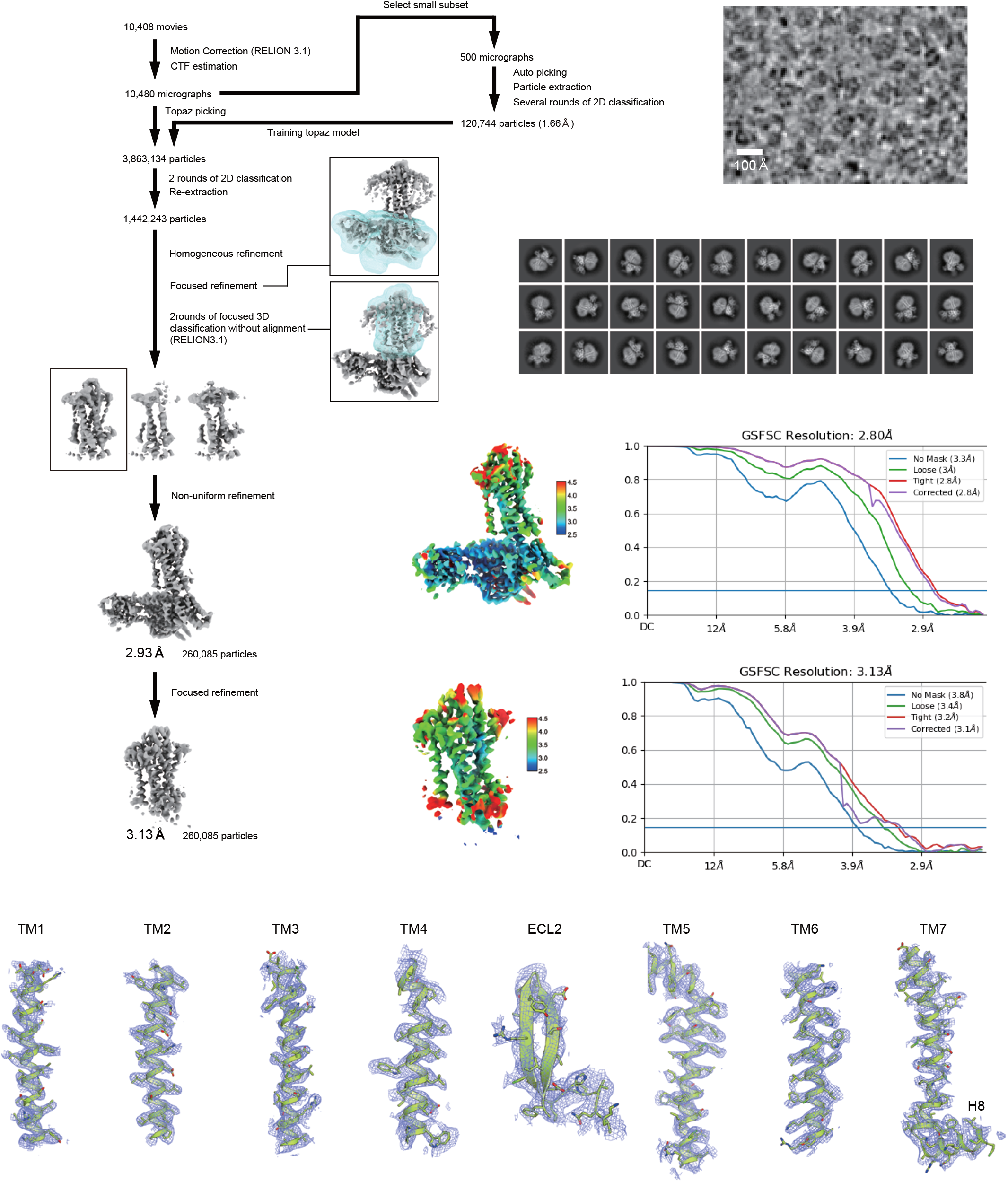
Cryo-EM workflow, maps, and model quality. Flow chart of the cryo-EM data processing for the ET_B_ -G_i_ complex, including particle projection selection, classification, and 3D density map reconstruction. Local resolution maps, FSC curves, and cryo-EM density maps are also shown. Unless otherwise noted, analysis jobs were run on cryoSPARC v3.3. Details are provided in the Methods section.

**Figure 3 – figure supplement 1.**
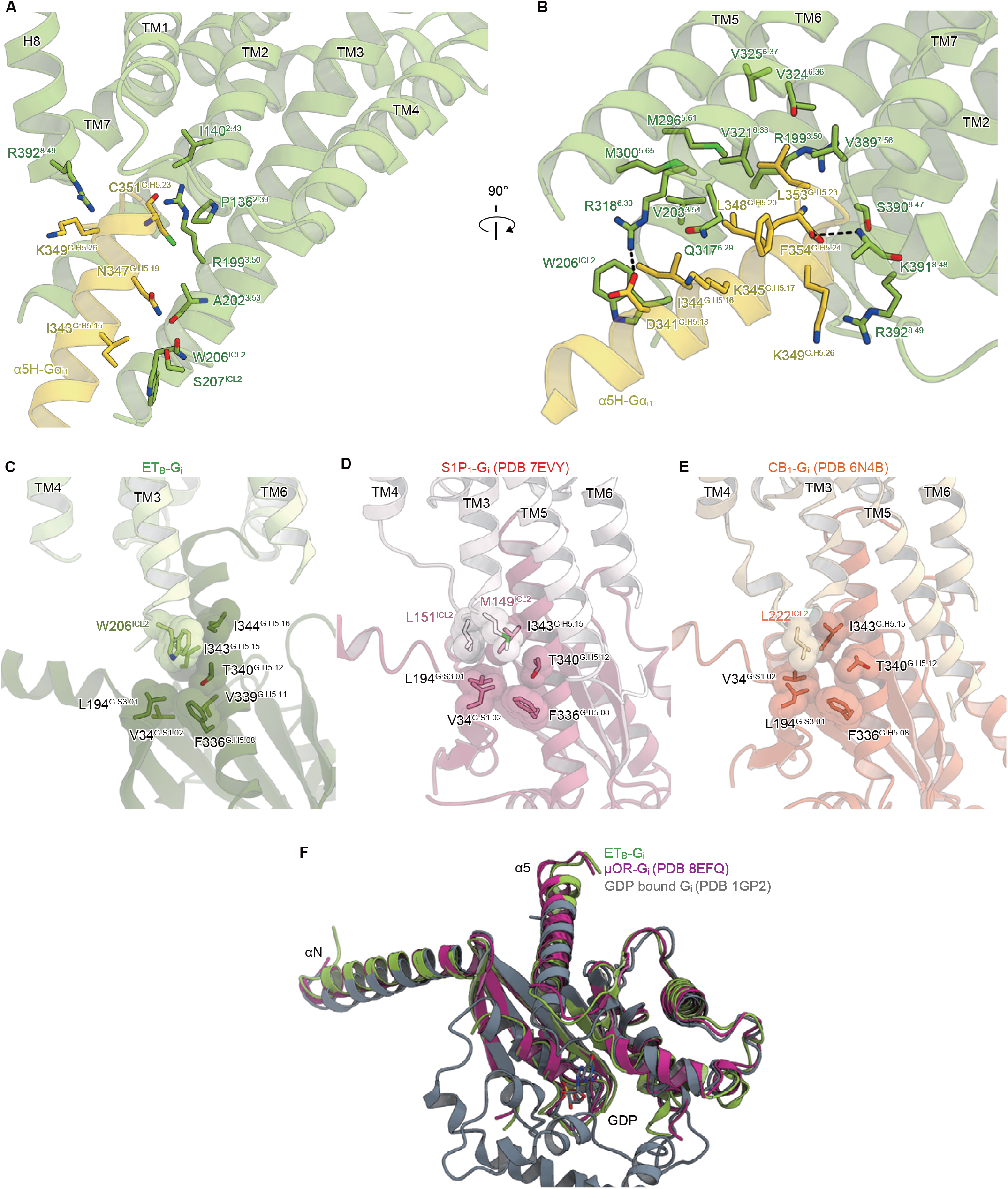
Detailed ET_B_-G_i_ interface. (A, B) Receptor-G_i_ interactions within 4.5 Å. Black dashed lines indicate hydrogen bonds. (C-E) Structural comparisons of the interactions between ICL2 and G_i_ in ET_B_ (C), S1P_1_ (D), and CB_1_ (E). Residues are shown as stick and CPK models. (F) Structural comparison of the Gα_i1_ subunits.

**Figure 3 – figure supplement 2.**
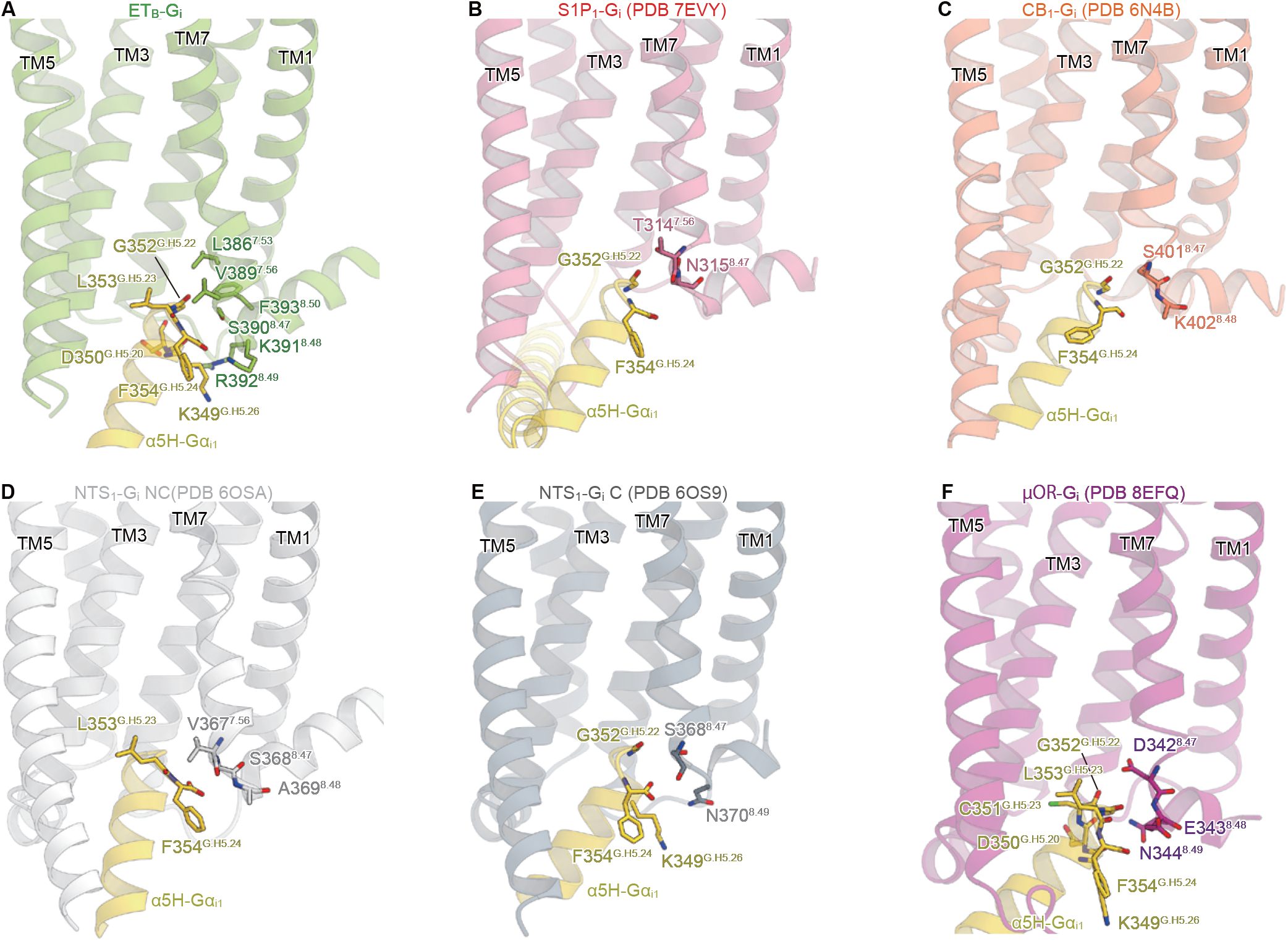
Comparison of the interactions between the α5-helix and TM7-H8. (A-F) Interactions between the α5-helix and TM7-H8 within 4.5 Å in the respective complexes.

## Notes

### Competing Interest Statement

O.N. is a co-founder and scientific advisor for Curreio.

## References

1. Yanagisawa, M. et al. A novel potent vasoconstrictor peptide produced by vascular endothelial cells. Nature 332, 411–415 (1988).

2. Maguire, J. J. & Davenport, A. P. Endothelin@25 - new agonists, antagonists, inhibitors and emerging research frontiers: IUPHAR Review 12. Br. J. Pharmacol. 171, 5555–5572 (2014).

3. Davenport, A. P. et al. Endothelin. Pharmacol. Rev. 68, 357–418 (2016).

4. Arai, H., Hori, S., Aramori, I., Ohkubo, H. & Nakanishi, S. Cloning and expression of a cDNA encoding an endothelin receptor. Nature 348, 730–732 (1990).

5. Sakurai, T. et al. Cloning of a cDNA encoding a non-isopeptide-selective subtype of the endothelin receptor. Nature 348, 732–735 (1990).

6. Haryono, A., Ramadhiani, R., Ryanto, G. R. T. & Emoto, N. Endothelin and the Cardiovascular System: The Long Journey and Where We Are Going. Biology 11, 759 (2022).

7. Barton, M. & Yanagisawa, M. Endothelin: 30 Years From Discovery to Therapy. Hypertens. Dallas Tex 1979 74, 1232–1265 (2019).

8. Doi, T., Sugimoto, H., Arimoto, I., Hiroaki, Y. & Fujiyoshi, Y. Interactions of endothelin receptor subtypes A and B with Gi, Go, and Gq in reconstituted phospholipid vesicles. Biochemistry 38, 3090–3099 (1999).

9. Inoue, A. et al. Illuminating G-Protein-Coupling Selectivity of GPCRs. Cell 177, 1933–1947.e25 (2019).

10. Tencé, M., Ezan, P., Amigou, E. & Giaume, C. Increased interaction of connexin43 with zonula occludens-1 during inhibition of gap junctions by G protein-coupled receptor agonists. Cell. Signal. 24, 86–98 (2012).

11. Koyama, Y. & Baba, A. Endothelin-induced cytoskeletal actin re-organization in cultured astrocytes: inhibition by C3 ADP-ribosyltransferase. Glia 16, 342–350 (1996).

12. Koyama, Y. Endothelin ETB Receptor-Mediated Astrocytic Activation: Pathological Roles in Brain Disorders. Int. J. Mol. Sci. 22, 4333 (2021).

13. Takai, M. et al. A potent and specific agonist, Suc-[Glu9,Ala11,15]-endothelin-1(8-21), IRL 1620, for the ETB receptor. Biochem. Biophys. Res. Commun. 184, 953–959 (1992).

14. Shihoya, W. et al. Activation mechanism of endothelin ET B receptor by endothelin-1. Nature 537, 363–368 (2016).

15. Nagiri, C. et al. Crystal structure of human endothelin ETB receptor in complex with peptide inverse agonist IRL2500. Commun. Biol. 2, 236 (2019).

16. Izume, T., Miyauchi, H., Shihoya, W. & Nureki, O. Crystal structure of human endothelin ETB receptor in complex with sarafotoxin S6b. Biochem. Biophys. Res. Commun. 528, 383–388 (2020).

17. Shihoya, W. et al. Crystal structures of human ETB receptor provide mechanistic insight into receptor activation and partial activation. Nat. Commun. 9, 4711 (2018).

18. Shihoya, W. et al. X-ray structures of endothelin ETB receptor bound to clinical antagonist bosentan and its analog. Nat. Struct. Mol. Biol. 24, 758–764 (2017).

19. Okuta, A., Tani, K., Nishimura, S., Fujiyoshi, Y. & Doi, T. Thermostabilization of the Human Endothelin Type B Receptor. J. Mol. Biol. 428, 2265–2274 (2016).

20. Ballesteros, J. A. & Weinstein, H. [19] Integrated methods for the construction of three-dimensional models and computational probing of structure-function relations in G protein-coupled receptors. in Methods in Neurosciences (ed. Sealfon, S. C.) vol. 25 366–428 (Academic Press, 1995).

21. Venkatakrishnan, A. J. et al. Molecular signatures of G-protein-coupled receptors. Nature 494, 185–194 (2013).

22. Duan, J. et al. Cryo-EM structure of an activated VIP1 receptor-G protein complex revealed by a NanoBiT tethering strategy. Nat. Commun. 11, 4121 (2020).

23. Dixon, A. S. et al. NanoLuc Complementation Reporter Optimized for Accurate Measurement of Protein Interactions in Cells. ACS Chem. Biol. 11, 400–408 (2016).

24. Xia, R. et al. Cryo-EM structure of the human histamine H1 receptor/Gq complex. Nat. Commun. 12, 2086 (2021).

25. Kim, K. et al. Structure of a Hallucinogen-Activated Gq-Coupled 5-HT2A Serotonin Receptor. Cell 182, 1574–1588.e19 (2020).

26. Nureki, I. et al. Cryo-EM structures of the β3 adrenergic receptor bound to solabegron and isoproterenol. Biochem. Biophys. Res. Commun. 611, 158–164 (2022).

27. Hattori, M., Hibbs, R. E. & Gouaux, E. A fluorescence-detection size-exclusion chromatography-based thermostability assay for membrane protein precrystallization screening. Struct. Lond. Engl. 1993 20, 1293–1299 (2012).

28. Akasaka, H. et al. Structure of the active Gi-coupled human lysophosphatidic acid receptor 1 complexed with a potent agonist. Nat. Commun. 13, 5417 (2022).

29. Liu, S. et al. Differential activation mechanisms of lipid GPCRs by lysophosphatidic acid and sphingosine 1-phosphate. Nat. Commun. 13, 731 (2022).

30. Yuan, Y. et al. Structures of signaling complexes of lipid receptors S1PR1 and S1PR5 reveal mechanisms of activation and drug recognition. Cell Res. 31, 1263–1274 (2021).

31. Xu, Z. et al. Structural basis of sphingosine-1-phosphate receptor 1 activation and biased agonism. Nat. Chem. Biol. 18, 281–288 (2022).

32. Zhuang, Y. et al. Molecular recognition of morphine and fentanyl by the human μ- opioid receptor. Cell 185, 4361–4375.e19 (2022).

33. Hua, T. et al. Activation and Signaling Mechanism Revealed by Cannabinoid Receptor-Gi Complex Structures. Cell 180, 655–665.e18 (2020).

34. Rasmussen, S. G. F. et al. Crystal structure of the β2 adrenergic receptor-Gs protein complex. Nature 477, 549–555 (2011).

35. Ring, A. M. et al. Adrenaline-activated structure of β2-adrenoceptor stabilized by an engineered nanobody. Nature 502, 575–579 (2013).

36. Cherezov, V. et al. High-resolution crystal structure of an engineered human beta2-adrenergic G protein-coupled receptor. Science 318, 1258–1265 (2007).

37. Manglik, A. et al. Crystal structure of the μ-opioid receptor bound to a morphinan antagonist. Nature 485, 321–326 (2012).

38. Huang, W. et al. Structural insights into μ-opioid receptor activation. Nature 524, 315–321 (2015).

39. Carpenter, B. & Tate, C. G. Active state structures of G protein-coupled receptors highlight the similarities and differences in the G protein and arrestin coupling interfaces. Curr. Opin. Struct. Biol. 45, 124–132 (2017).

40. Deupi, X. et al. Stabilized G protein binding site in the structure of constitutively active metarhodopsin-II. Proc. Natl. Acad. Sci. U. S. A. 109, 119–124 (2012).

41. Goncalves, J. A. et al. Highly conserved tyrosine stabilizes the active state of rhodopsin. Proc. Natl. Acad. Sci. U. S. A. 107, 19861–19866 (2010).

42. Flock, T. et al. Universal allosteric mechanism for Gα activation by GPCRs. Nature 524, 173–179 (2015).

43. Kato, H. E. et al. Conformational transitions of a neurotensin receptor 1–Gi1 complex. Nature 572, 80–85 (2019).

44. Wall, M. A. et al. The structure of the G protein heterotrimer Gi alpha 1 beta 1 gamma 2. Cell 83, 1047–1058 (1995).

45. Su, M. et al. Structural Basis of the Activation of Heterotrimeric Gs-Protein by Isoproterenol-Bound β1-Adrenergic Receptor. Mol. Cell 80, 59–71.e4 (2020).

46. Okamoto, H. H. et al. Cryo-EM structure of the human MT1-Gi signaling complex. Nat. Struct. Mol. Biol. 28, 694–701 (2021).

47. Du, Y. et al. Assembly of a GPCR-G Protein Complex. Cell 177, 1232–1242.e11 (2019).

48. Kobayashi, K. et al. Cryo-EM structure of the human PAC1 receptor coupled to an engineered heterotrimeric G protein. Nat. Struct. Mol. Biol. 27, 274–280 (2020).

49. Zivanov, J. et al. New tools for automated high-resolution cryo-EM structure determination in RELION-3. eLife 7, (2018).

50. Punjani, A., Rubinstein, J. L., Fleet, D. J. & Brubaker, M. A. cryoSPARC: algorithms for rapid unsupervised cryo-EM structure determination. Nat. Methods 14, 290–296 (2017).

51. Bepler, T. et al. Positive-unlabeled convolutional neural networks for particle picking in cryo-electron micrographs. Nat. Methods 16, 1153–1160 (2019).

52. Emsley, P., Lohkamp, B., Scott, W. G. & Cowtan, K. Features and development of Coot. Acta Crystallogr. D Biol. Crystallogr. 66, 486–501 (2010).

53. Adams, P. D. et al. PHENIX: a comprehensive Python-based system for macromolecular structure solution. Acta Crystallogr. D Biol. Crystallogr. 66, 213–221 (2010).

54. Afonine, P. V. et al. Real-space refinement in PHENIX for cryo-EM and crystallography. Acta Crystallogr. Sect. Struct. Biol. 74, 531–544 (2018).

